# Prokaryotic rRNA-mRNA interactions are involved in all translation steps and shape bacterial transcripts

**DOI:** 10.1101/2020.07.24.220731

**Authors:** Shir Bahiri Elitzur, Rachel Cohen-Kupiec, Dana Yacobi, Larissa Fine, Boaz Apt, Alon Diament, Tamir Tuller

## Abstract

The well-established Shine-Dalgarno model suggests that translation initiation in bacteria is regulated via base-pairing between ribosomal RNA (rRNA) and mRNA. However, little is currently known about the contribution of such interactions to the rest of the translation process and to the way bacterial transcript evolve. We used novel computational analyses and modelling of 823 bacterial genomes coupled with experiments to demonstrate that rRNA-mRNA interactions are diverse and regulate not only initiation, but all translation steps from pre-initiation to termination across the many bacterial phyla that have the Shine-Dalgarno sequence. As these interactions dictate translation efficiency, they serve as a driving evolutionary force for shaping transcripts in bacteria. We observed selection for strong rRNA-mRNA interactions in regions where such interactions are likely to enhance initiation, regulate early elongation and ensure the fidelity of translation termination. We discovered selection *against* strong interactions and *for* intermediate interactions in coding regions and present evidence that these interactions maximize elongation efficiency while also enhancing initiation by ‘guiding’ free ribosomal units to the start codon.

**Importance:** Previous research has reported the significant influence of rRNA-mRNA interactions mainly in the initiation phase of translation. The results reported in this paper suggest that, in addition to the rRNA-mRNA interactions near the start codon that trigger initiation in bacteria, rRNA-mRNA interactions affect all sub-stages of the translation process (pre-initiation, initiation, elongation, termination). In addition, these interactions affect the way evolutionary forces shape the bacterial transcripts while considering trade-offs between the effects of different interactions across different transcript regions on translation efficacy and efficiency. Due to the centrality of the translation process, these findings are relevant to all biomedical disciplines.

## Introduction

The Shine-Dalgarno (SD) sequence or ribosome binding site (RBS) region, which is approximately 8-10 nucleotides upstream of the start codon in prokaryotic mRNA(1, 2) is known to be involved in prokaryotic translation initiation via base-pairing to a complementary sequence in the 16S rRNA component of the small ribosomal subunit – the anti Shine-Dalgarno (aSD)(1–8).

Recent studies have also suggested that sequences (motifs) within the coding region that interact with the aSD, similarly to the SD, can slow down or pause translation elongation in several bacteria species(9–12). Thus, it was suggested that such sequences in the coding region decrease the overall translation rate, and can generally be considered deleterious. Other studies have suggested that selection against internal SD-like sequences which promote rRNA-mRNA interactions can act against codons that tend to compose such motifs(13–15).

Here, based on a comprehensive analysis of 823 prokaryotic genomes we provide a high resolution model of rRNA-mRNA interactions during translation, which suggest that the current knowledge about the function of rRNA-mRNA interactions is just the ‘tip of the iceberg’: in most of the bacteria analyzed, rRNA-mRNA interactions seem to be involved in all stages and sub-stages of translation and not just in the initiation phase as was known to date (see Supplementary section S.1 and Figure S1). Thus, rRNA-mRNA interactions affect the way evolution shapes the nucleotide composition along the entire transcript to optimize translation.

## Results

To understand the interactions between the 16S rRNA and mRNAs across the bacterial kingdom, we developed a high-resolution computational model to predict the strength of rRNA-mRNA interactions, where low hybridization free energy indicates a stronger interaction (Material and Methods section). We used our model to analyze the entire transcriptome of 823 bacterial species, investigating all possible positions across all transcripts (i.e. 2,896,245 transcripts). To detect patterns of evolutionary selection, we compared the distribution of rRNA-mRNA interaction strength in each position along the transcriptome of each genome to the one expected by a null model. The null model preserves the codon frequencies, amino acid content, and GC content in each transcript (Methods section).

For each position along the transcriptome we performed three statistical tests to answer the following questions:

1) Does the nucleotide (nt) sequences in that position tend to produce *stronger* rRNA-mRNA interactions than expected by the null model?

A positive answer to this question supports the conjecture that strong interactions in the position in the 5’ UTR, the beginning and end of the coding region tend to improve translation.

2) Does the nt sequences in that position tend to produce *weaker* rRNA-mRNA interactions than expected by the null model?

A positive answer to this question supports the conjecture that weak interactions in the position in the coding region tend to improve translation.

3) Does the the nt sequences in that position tend to produce *intermediate* (moderate strength: neither very strong nor very weak) rRNA-mRNA interactions in comparison to what is expected by a null model? (see Figure 1A and Methods).

**Figure 1.**
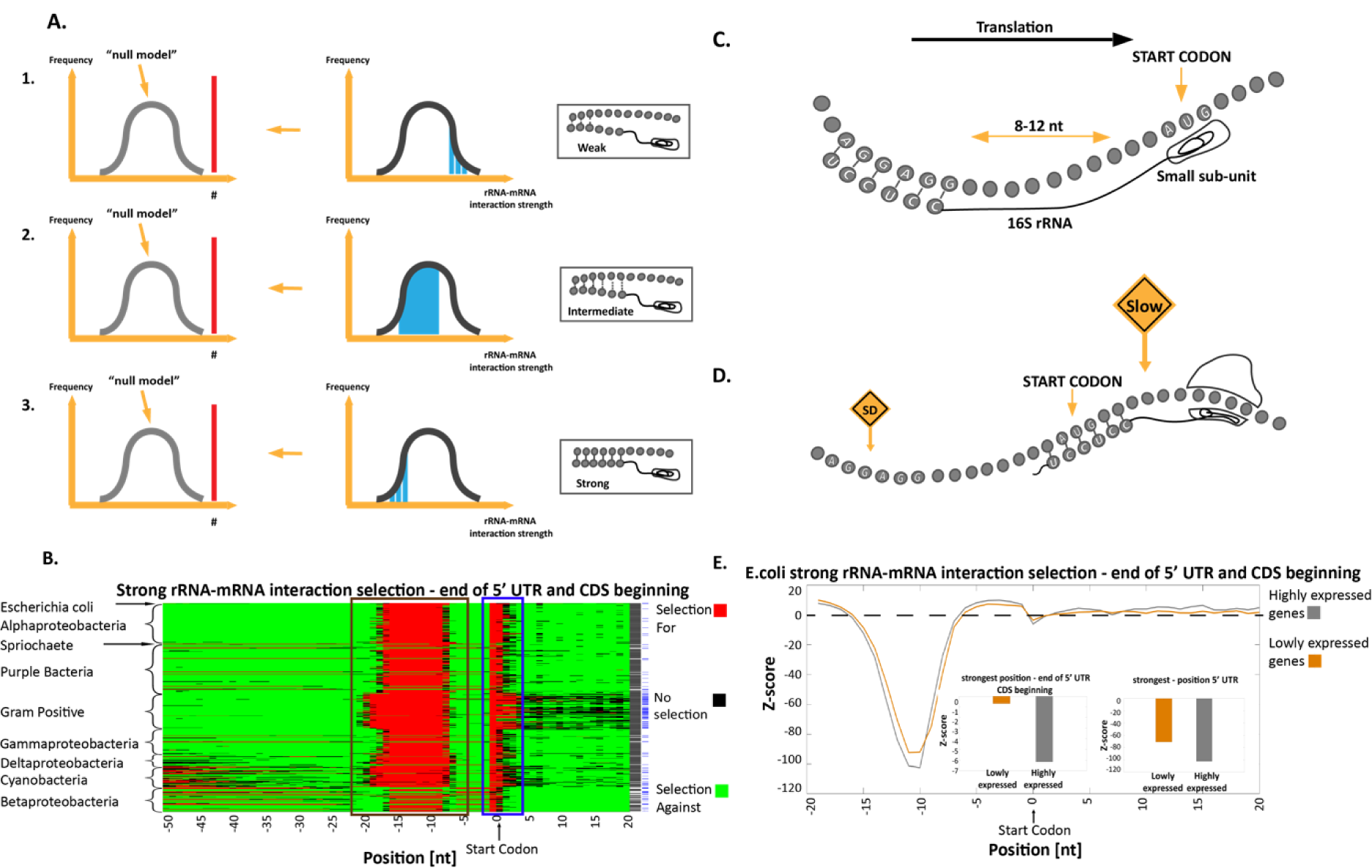
Prediction of rRNA-mRNA interaction strength and selection for or against strong rRNA-mRNA interactions at the 5’UTR and at the beginning of the coding region. **A**. The three statistical tests to detect evolutionary selection for different rRNA-mRNA interaction strength (see Methods). 1. Enrichment of sub-sequences with weak rRNA-mRNA interactions; 2. intermediate rRNA-mRNA interactions, and 3. strong rRNA-mRNA interactions. We examined weak, intermediate and strong rRNA-mRNA interaction strengths separately, and in each case tested if their number or mean value was significantly higher than expected by the null model. **B**. Results of the test for selection of strong interactions in the 5’UTR and first 20 nucleotides of the coding region. Each row represents a bacterium, rows are clustered based on phyla, and each column is a position in the transcripts of the analyzed organisms. Red and green indicate a position with significant selection for and against strong rRNA-mRNA interaction in comparison to the null model, respectively. Black indicates a position with no significant selection (Material and Methods section). Second column from the right: a black pixel represents a bacterium for which the number of positions with significant selection for strong interactions was significantly higher than the null model in the 5’UTR. Rightmost column: a blue pixel represents a bacterium for which the number of significant positions with selection for strong interactions was significantly higher than the null model in the last nucleotide of the 5’UTR and the first 5 nucleotides of the coding region. **C**. An illustration of the way strong rRNA-mRNA interactions affect translation initiation: The rRNA-mRNA interactions upstream of the start codon initiate translation by aligning the small subunit of the ribosome to the canonical start codon. **D**. An illustration of the suggested model: strong interactions at the first steps of elongation slow down the ribosome movement. **E**. Z-scores for rRNA-mRNA interaction strengths at the last 20 nucleotides of the 5’UTR and first 20 nucleotides of the coding regions in highly and lowly expressed E. coli genes. Lower/higher Z-scores indicate selection for/against strong rRNA-mRNA interactions respectively, in comparison to what is expected by the null model. Highly and lowly expressed genes were selected according to protein abundance. Insets: two bar graphs of the Z-scores in highly and lowly expressed genes in the two regions of the reported signals.

A positive answer to this question supports the conjecture that intermediate interactions in the position in the coding region tend to improve translation.

We report the observed tendencies of sub-sequences within different transcript regions to produce strong, intermediate, and weak interactions with the 16S rRNA.

### Selection for strong rRNA-mRNA interactions at the 5’UTR end and at the beginning of the coding region to regulate translation initiation and early translation elongation

First, we analyzed the 5’UTRs of 551 bacteria with aSD (anti Shine Delgarno) sequence in the rRNA. It was suggested that translation initiation in prokaryotes is initiated by hybridization of the 16S rRNA to the mRNA(2). The 16S rRNA binds to the 5’UTR near and upstream of the START codon(4) as depicted in Figure 1C. Indeed, as can be seen in Figure 1B (brown box) in almost all of the analyzed bacteria, there is a significant signal of selection for *strong* rRNA-mRNA interactions at positions -8 through -17 relative to the START codon, in agreement with the Shine-Dalgarno model(1, 2).

A second signal of selection for *strong* rRNA-mRNA interactions appears in the last nucleotide of the 5’UTR and the first five nucleotides of the coding sequence (Figure 1B, blue box). Since the elongating ribosome is positioned around 11 nucleotides downstream of the position its rRNA interacts with the mRNA (16), it is likely that these rRNA-mRNA interactions are related to slowing down the early elongation phase of the ribosome.

It has been suggested that at the beginning of the coding region there are various features that slow down the early stages of translation elongation to improve organism fitness, e.g. via optimizing ribosomal allocation and chaperon recruitment (Figure 1D)(17, 18). It is likely that this second novel signal is a mechanism of such regulation. Both of the reported signals above occur in 89% of the analyzed bacteria.

A comparison of highly and lowly expressed genes in *E. coli* (Figure 1E) reveals that both signals are stronger in the highly expressed genes, which are under stronger selection to optimize translation. The difference between the Z-scores of highly and lowly expressed genes in the two reported signal regions was highly significant (nucleotides -8 through -17 in the 5’UTR: Wilcoxon rank-sum test p=7.9·10^−5^; last nucleotide of the 5’UTR and the first 5 nucleotides of the coding sequence: Wilcoxon rank-sum test p=9.3·10^−4^).

### Selection against strong rRNA-mRNA interactions in the coding regions that prevents the slowing down of translation elongation

Ribo-seq analyses in *E. coli* have indicated that strong interactions between the 16S rRNA and the mRNA can lead to pauses during translation elongation, hindering translation(9–12, 19) (Figure 2D). Avoiding such strong rRNA-mRNA interactions in the coding region should thus allow the ribosome to flow efficiently during translation elongation. The deleterious effects of such strong rRNA-mRNA interaction sequences may also be due to their role in encouraging internal translation initiation which would create truncated and frame-shifted protein products. Hockenberry et al(15) found support for this claim by observing that the occurrence of AUG start codons is significantly depleted downstream of existing strong rRNA-mRNA interaction sequences in *E. coli*.

**Figure 2.**
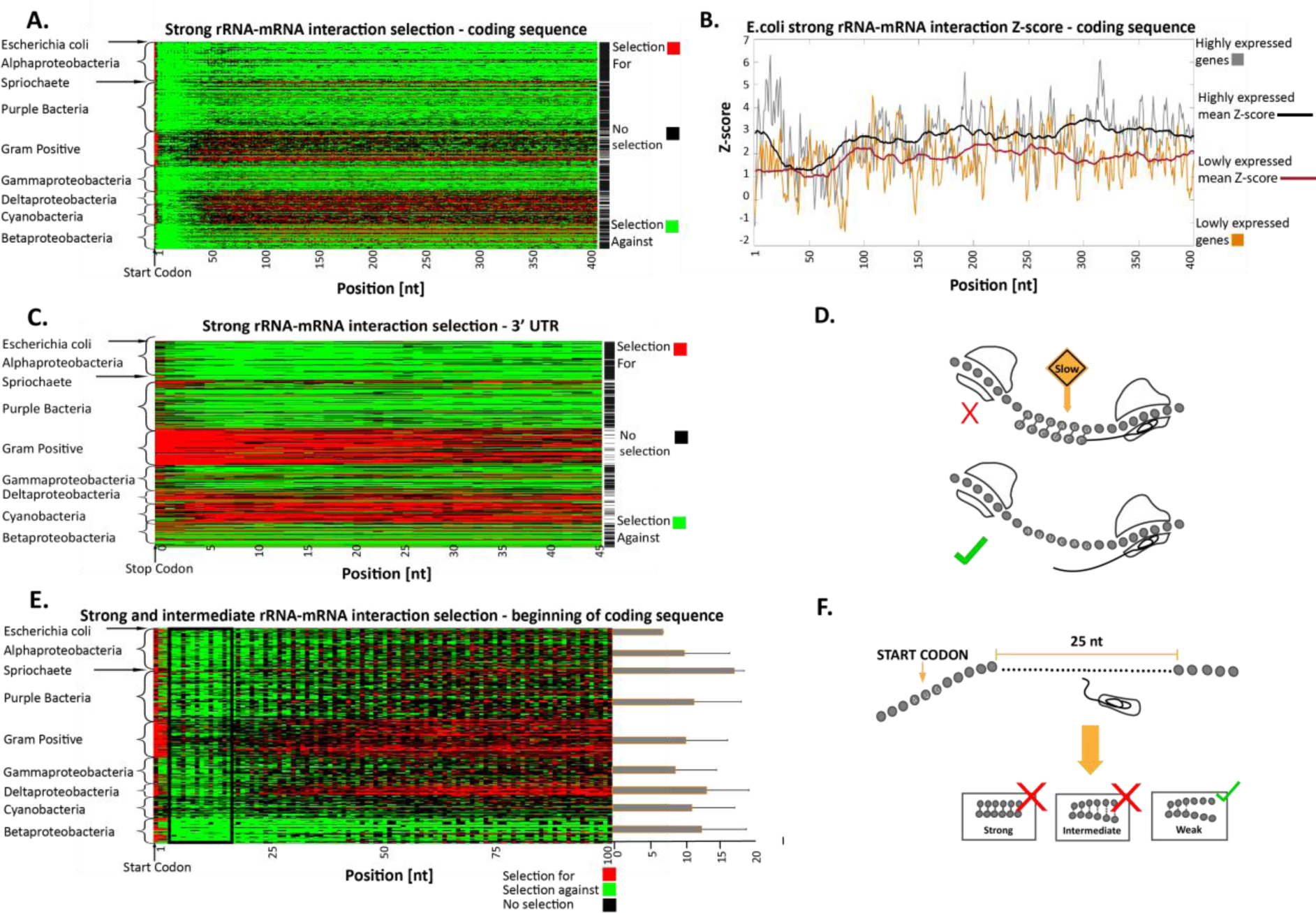
Selection for and against strong rRNA-mRNA interactions in the coding and 3’UTR regions. **A**. The positions with selection for or against strong rRNA-mRNA interaction in the first 400 nt of coding regions. Each row represents a bacterium, the rows clustered by phyla, and each column is a position in the transcripts of the analyzed organisms. Red/green indicates a position with significant selection for/against strong rRNA-mRNA interactions in comparison to the null model, respectively (Material and Methods section). Black indicates positions with no significant selection. Rightmost column: black represents bacteria for which the number of positions with significant selection against strong interactions was significantly higher than the null model. **B**. Z-score for rRNA-mRNA interaction strength at the first 400 nucleotides of the coding regions in highly and lowly expressed genes in E. coli. Lower/higher Z-scores mean stronger/weaker rRNA-mRNA interactions than the null model, respectively. The bold black/red lines represent a 40-nucleotide moving average in highly/lowly expressed genes, respectively. **C**. Positions with selection for or against strong rRNA-mRNA interaction strength in the 3’ UTR. Each row represents a bacterium, the rows are clusteredby phyla, and each column is a position in the bacteria’s transcript. Red/green indicates a position with significant selection for/against strong rRNA-mRNA interactions relative to the null model, respectively (Material and Methods section). Black indicates position with no significant selection. Rightmost column: black represents bacteria for which the number of significant positions with selection against strong interactions is significantly higher than in the null model. **D**. The effect of strong rRNA-mRNA interactions in the coding region on translation elongation: such interactions can slow down ribosome movement and retard translation. **E**. Positions with significant strong and intermediate rRNA-mRNA interaction distribution in the first 100 nt of the coding region. Each row represents a bacterium, the rows are clustered by phyla, and each column is a transcript position. Red/green indicates a position with significant selection for/against strong and intermediate rRNA-mRNA interactions, in comparison to the null model, respectively (Material and Methods section). Black indicates position with no significant selection. Bars at right of plot show: for each bacterium, we calculated in a sliding window of 40 nucleotides the number of positions with selection against strong and intermediate interactions. The bars represent the average number of windows at the beginning of the coding region that had more selection against strong and intermediate interactions than the rest of the transcript, averaged by phylum. Lines extending from bars represent standard deviations (the periodicity in the signal is related to the genetic code). **F**. An illustration of our model: strong and intermediate interactions at the first 25 nucleotides can be deleterious and can promote initiation from erroneous positions.

Our analysis reveals evidence of significant selection against *strong* rRNA-mRNA interactions in the coding region (Figure 2A). In 55% of the bacteria analyzed, at least 50% of the positions in the first 400 nucleotides of the coding region exhibit a signal of significant selection against *strong* rRNA-mRNA interactions. Importantly, this selection was also observed away from positions that are upstream of a nearby AUG, suggesting that such selection is also related to elongation, and not just to avoiding internal translation initiation (Figure S2 and Supplementary section S.2). Our findings are in agreement with those of Yang et al(19) who showed depletion in internal-SD-like sequences in most species analyzed (without a control for the positions that are close to an AUG). However, this study provides this insight at a much higher resolution: Yang et al examined the occurrence of such sequences over the total genome whereas we performed a per-position comparison in each genome.

We found evidence for selection against *strong* rRNA-mRNA interactions in the coding region throughout the bacteria phyla analyzed, except for in cyanobacteria and gram positive bacteria which seem to exhibit selection for *strong* rRNA-mRNA interactions (Figure 2A). It has been hypothesized that interactions between rRNA and mRNA are weaker in cyanobacteria as 16S ribosomal RNA is folded in such a way that subsequences that usually interact with the mRNA are situated within the RNA structure(20, 21). Thus, in these organisms, it is expected that rRNA-mRNA interactions are less probable, resulting in lower selection pressure to eliminate sub-sequences that can interact with the rRNA in the coding region. A similar trend can be seen in the 3’UTR of genes (Figure 2C). We postulate that similar to cyanobacteria, gram positive bacteria also have rRNA structures that result in less efficient rRNA-mRNA interactions.

Again, a comparison between highly and lowly expressed genes in *E. coli* reveals that selection against nucleotide sequences leading to strong interactions in the coding region is stronger for highly expressed genes which are under stronger selective pressure for more accurate and efficient translation (Wilcoxon rank-sum test p=1.5·10^−30^; Figure 2B).

In addition, as can be seen in Figure 2E: At the beginning of the coding region (5-25 nucleotides), there is significant increased selection against *strong* and *intermediate* rRNA–mRNA interactions (typical p-value 0.0097). The presence of sub-sequences that interact in a strong/intermediate manner near the beginning of the coding region is probably more deleterious as it might promote with higher probability initiation from erroneous positions (see illustration in Figure 2F); indeed, similar signals related to eukaryotic and prokaryotic initiation were reported (13, 22).

### Selection for strong rRNA-mRNA interactions at the end of the coding sequences to improve the fidelity of translation termination

In 82% of the analyzed bacterial species, in 50% of the positions at the last 20 nucleotides of the coding region, there is selection for *strong* rRNA-mRNA interactions (Figure 3A). It is likely that this constitutes a mechanism for slowing ribosome movement when approaching the stop codon, and serves to ensure efficient and accurate termination and prevent translation read-through(23, 24) (Figure 3F). Researchers have suggested that this selection may have the function of assisting initiation of overlapping or nearby downstream genes in operons(13); however, we observed this phenomenon universal across all genes and bacteria, including the last genes in an operon which are not closely followed by other genes (Supplementary section S.3 and Figure S3).

**Figure 3.**
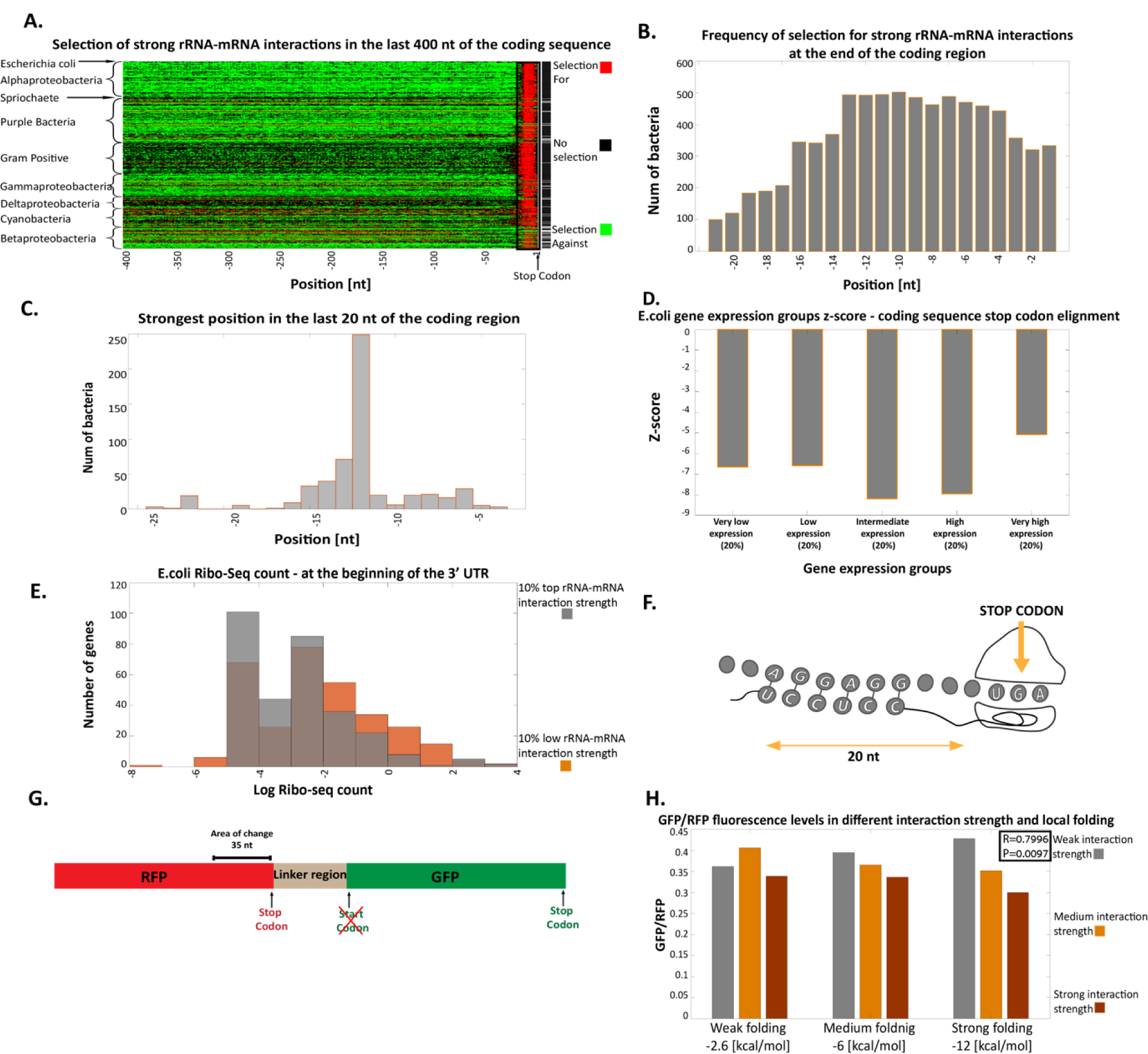
Selection for/against strong rRNA-mRNA interactions at the end of the coding region. **A**. Selection for or against strong rRNA-mRNA interaction in the last 400 nt of the coding regions. Each row represents a bacterium, rows are clustered by phyla, and each column is a position in the bacterial transcript. Red/green indicates positions with significant selection for/against strong rRNA-mRNA interaction in comparison to the null model, respectively (Material and Methods section). Black indicates positions with no significant selection. Rightmost column: black pixels represent bacteria where the number of significant positions with selection for strong interactions was significantly higher than the null model. **B**. The number of bacteria with significant selection for strong rRNA-mRNA interactions in each of the last 20 nt of the coding region. **C**. Distribution of the position with the lowest rRNA-mRNA interaction Z-score, indicating the strongest rRNA-mRNA interaction, in the last 20 nt of the coding region among the analyzed bacteria. **D**. Mean of the lowest Z-score for rRNA-mRNA interaction strength among the last 20 nucleotides of the coding region for groups of genes classified according to gene expression levels. **E**. Ribo-seq analysis: average Ribo-seq read count distribution at the beginning of the 3’UTR for genes with strong (gray bars) vs. weak (orange bars) rRNA-mRNA interactions at the end of the coding sequence (Material and Methods section). **F**. An illustration of our model: strong interactions at the end of the coding region enhance the accurate recognition of the stop codon and aid in translation termination. **G**. The experiment construct, an RFP gene connected to a GFP gene. We tested the effect of different rRNA-mRNA interaction strengths in the last 35 nt of the RFP gene by creating variants with different folding in the last 40 nt. **H**. Bar graph of values proportional to GFP / RFP fluorescence levels in the 9 variants (see Methods) grouped according to their local folding energies.

It has previously been found that when the rRNA binds to the mRNA the ribosome is generally decoding a codon located approximately 11 nt downstream of the binding site(9). To validate this, we inferred the positions with selection for the strongest interactions and identified those with minimum rRNA-mRNA interaction Z-scores within the last 20 nt of the coding region, in most of the analyzed bacteria (Material and Methods section). We discovered that the strongest and most significant positions across all bacteria are indeed -9 through -22 relative to the STOP codon (Figures 3B and 3C). This supports our hypothesis that the interactions indeed function to halt the ribosome on the STOP codon and not to initiate the next open reading frame in the operon.

We examined the relationship between the strength of selection for strong interaction in the last 20 nt of coding regions with different levels of gene expression and found it to be convex: such selection is stronger for genes with intermediate expression and weaker for both lowly- and highly-expressed genes (Figure 3D). We consider that the weaker selection in lowly-expressed genes may be due to lower selection pressure on the gene in general(25). Conversely, the weaker signal in highly-expressed genes may be due to stronger selection on translation elongation and termination rates: the ribosome density in these genes is higher(17), and if a ribosome is stalled in order to promote accurate termination it may cause ribosome queuing at the 3’-end, resulting in inefficient ribosomal allocation. Highly expressed genes may have other mechanisms for ensuring termination fidelity.

To test if strong rRNA-mRNA interactions just prior to the stop codon improve termination fidelity, we analyzed Ribo-seq data of *E.coli*(26) (Figure 3E and Material and Methods section). We expected that if such an interaction improves the fidelity of termination, mRNAs with a strong interaction will exhibit less read-through events and thus we will observe less Ribo-seq read counts (RC) downstream of the STOP codon. Indeed, we found that the average read count for the 20 nucleotides after the stop codon was lower following genes with strong rRNA-mRNA interactions in the last 20 nucleotides of the coding region, compared to genes with weaker interactions in this region (mean RC=0.334 and 0.514, respectively; Wilcoxon rank-sum test p=0.001).

To further experimentally test our hypothesis of strong rRNA-mRNA interactions just prior to the stop codon preventing stop-codon read-through, we used a construct mRNA with a gene coding for red fluorescent protein (RFP) linked to a gene coding for green fluorescent protein (GFP; Figure 3G). We positioned the GFP gene downstream such that its expression acts as an indicator of read-through expression, and variants with higher GFP fluorescence are indicative of higher rates of stop-codon read-through (Material and Methods section and Supplementary S.4 and Figure S4). We designed nine variants with different rRNA-mRNA interaction strengths and local mRNA folding at the last 40 nt(27) of the RFP, and measured their florescence. As hypothesized, we found that variants with stronger rRNA-mRNA interactions at the end of the RFP coding region tend to produced lower levels of GFP (Figure 3H). We found that there is high correlation between the relative read-through signal (the ration between the GFP fluorescence and the RFP florescence) and the predicted rRNA-mRNA interactions strength prior to the stop codon even when controlling for the local mRNA folding near the stop codon (partial Spearman correlation: r=0.6997 P=0.0096).

### Selection for intermediate rRNA-mRNA interactions in the coding region and UTRs to improve the pre-initiation diffusion of the small subunit to the initiation site

The previous sections presented evidence for selection against strong interactions between the rRNA and mRNA throughout most of the coding region, but this doesn’t mean that all interactions throughout this region are deleterious: other forces may act in differing directions. Prior to binding with mRNA, free ribosomal units travel by diffusion. Some interaction with the mRNA may assist to ‘guide’ the diffusing small subunit of the ribosome to remain near the transcript and ‘help’ them find the start codon, increasing their diffusion efficiency and consequently overall translation initiation efficiency (Figure 4F.1).

**Figure 4.**
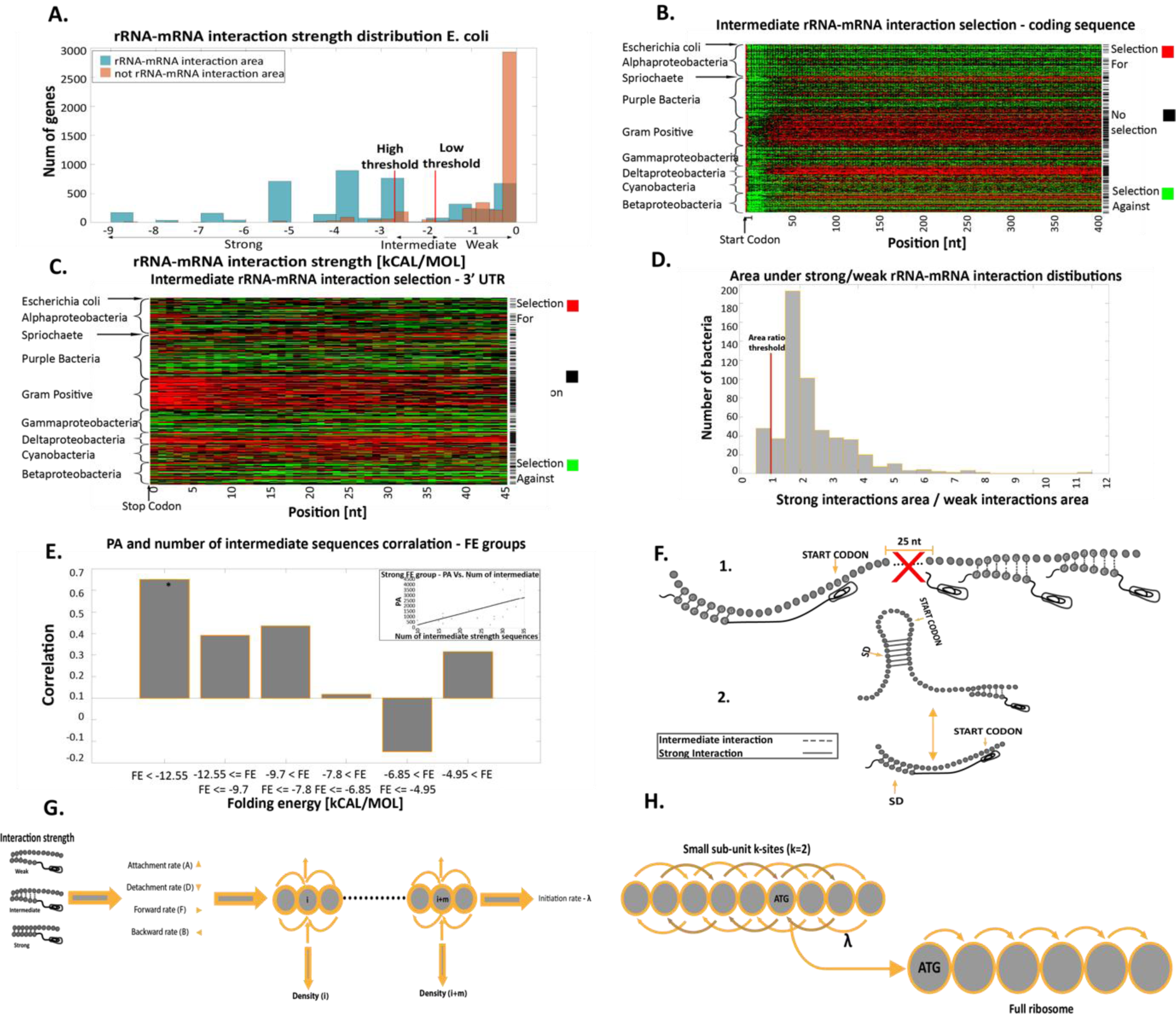
Selection for/against intermediate rRNA-mRNA interactions in the coding and UTR regions. **A**. Definition and threshold validation for intermediate-strength rRNA-mRNA interactions in E. coli. Two distributions are shown: 1. blue bars: maximum rRNA-mRNA interaction strength distribution of the interaction strength region related to region 1 (see main text). 2. Orange bars: maximum rRNA-mRNA interaction strength distribution in the weak interaction region (related to region 2) (see main text). Thresholds for defining intermediate interactions for this organism are also depicted. **B**. Positions with selection for high/low number of intermediate rRNA-mRNA interactions in first 400 nt of the coding regions. Rows represent individual bacteria and are clustered by phyla; each column is a transcript position. Red/green indicate positions with significant selection for/against intermediate rRNA-mRNA interaction relative to the null model, respectively (Material and Methods section). Black indicates positions with no significant selection. Rightmost column: black pixels represent bacteria where the number of positions with significant selection for intermediate interactions is significantly higher than the null model. **C**. Positions with selection for high/low number of intermediate rRNA-mRNA interactions in the 3’ UTR. Rows represent bacteria and are clustered by phyla; each column is a transcript position. Red/green indicates positions with significant selection for/against intermediate rRNA-mRNA interactions relative to the null model, respectively (Material and Methods section). Rightmost column: black pixels represent bacteria where the number of positions with significant selection for intermediate interaction is significantly higher than the null model. **D**. Distribution of the area ratio. A ratio larger than 1 suggests that it is more probable that the inferred thresholds are related to (intermediate) rRNA-mRNA interactions, and not to a lack of interaction. **E**. The number of intermediate sequences and PA correlations in GFP synonymous variants. The GFP variants are divided into six groups according to their FE near the start codon. The FE thresholds were selected so as to have approximately equal numbers of GFP variants in each group. Groups with significant correlation are marked with *. Inset: correlation between PA and number of intermediate interaction sequences for the strongest FE group. **F**. Illustration of intermediate interaction effects on translation initiation. 1) Intermediate interactions in the coding sequence. 2) This aid initiation when there is strong mRNA folding in the region surrounding the START codon (i.e. when initiation is more rate limiting). **G**. An illustration of the biophysical model. Each site’s parameters are determined by its rRNA-mRNA interaction strength. There is an attachment rate to the site, detachment rate from the site, movement forward to the site and from it and movement backward from the site and to it. This model allows for deduction of the initiation rate for insertion into the elongation model. **H**. An illustration of the rRNA-mRNA interaction strength extended model. The density of each site is determined by k sites before it and k sites after it. (Supplementary section S.8).

Initiation is often the rate limiting stage of translation and the most limiting aspects probably appear to be the diffusion of the small sub-unit to the SD region. Diffusion along the mRNA may be faster: if mRNAs can ‘catch’ small ribosomal sub-units and then direct them to their start codons, they may be favored by evolution. The large amount of redundancy in the genetic code allows for mutations that may improve interactions between the rRNA and mRNA even in the coding region, without negatively affecting protein products; however as we have seen, strong interactions in the coding region are problematic. Based on these considerations; we hypothesized that evolution shapes coding regions to include intermediate rRNA-mRNA interactions, which are not strong enough to halt elongation, but can optimize pre-initiation diffusion.

To test this hypothesis, we created an unsupervised optimization model to identify sequences with intermediate rRNA-mRNA interactions by adaptively calculating rRNA-mRNA interaction-strength thresholds for each bacterium. The algorithm selects rRNA-mRNA interaction strength thresholds such that they delineate the maximum number of significant positions with rRNA-mRNA interactions between these thresholds (see more details in the Material and Methods section).

To verify that the thresholds are reasonable, we looked at the highest (per gene) rRNA-mRNA interaction strength distribution in the 5’UTR in two regions: 1) The canonical rRNA-mRNA interaction region during initiation (i.e. nucleotides -8 through -17 upstream to the start codon). 2) The region in the 5’UTR which is upstream to 1). We then defined each gene by two values: *a*. Minimum interaction strength (i.e. strongest interaction) from region 1) distribution. *b*. Minimum interaction strength from region 2) distribution. For each bacterium, we created distribution plots based on values *a*. and *b*. over its genes. Figure 4A includes these two distributions for *E. coli*; as can be seen, the rRNA-mRNA intermediate interaction strength thresholds for this bacterium are in the overlapping region of the two distributions. Furthermore, we calculated the area between the optimized intermediate thresholds under the distribution of all values of rRNA-mRNA interaction strength in the aforementioned regions (1) and (2) (Figure 4D). As expected, the area under distribution 1) is greater than the area under distribution 2) in most of the bacteria (the ratio is larger than 1 in 91 percent of the bacteria). This provides confirmation that the range of interaction strengths identified corresponds to intermediate interactions and not to a lack of interaction.

Our analyses revealed that in 52% of the analyzed bacteria at least 50% of the positions are under significant selection for *intermediate* rRNA-mRNA interactions: according to the null model this would be expected to be the case for only 0.18% (Figure 4B). A similar trend can be seen in the 3’UTR (Figure 4C). The level of selection for intermediate interactions in the coding region varies among the bacterial Phylum and thus may be affected by various phylum-specific characteristics as growth rate, competition, and many aspects of translation regulation (Supplementary section S.5).

Our null model preserves the protein itself, the codon bias and the GC content. Therefore, the observed selection cannot be favouring specific codons or amino acids. In addition, our rRNA-mRNA interaction profiles consider all three reading frames; hence, the amino acids are not the key factor that influences this signal. Furthermore, the fact that we see a similar pattern of selection in the UTRs (Figure 4C) suggests that this pattern cannot be attributed only to selection for certain codon pairs.

We hypothesize that selection for intermediate rRNA-mRNA interactions in the coding region of a gene should improve its translation initiation efficiency and thus its protein levels. To demonstrate this, we calculated the partial Spearman correlations between the number of intermediate interaction sequences in the GFP variant (see previous section) and the heterologous protein abundance (PA), based on 146 synonymous GFP variants that were expressed from the same promoter(28). The control variables were the codon adaptation index (CAI) (29); a measure of codon usage bias, and mRNA folding energy (FE) near the start codon, known to affect translation initiation efficiency (the weaker the folding in the vicinity of the start codon the higher the fidelity and efficiency of translation initiation) (30–32).

We defined an area of intermediate interactions according to the thresholds determined by our model in *E*. coli and calculated the correlation explained above. As expected, the correlation was positive and significant (r=0.35; P=0.2·10^−4^) indicating that variants with more sub-sequences in the coding region that bind to the rRNA with an intermediate interaction strength tend to have higher PA.

We found that this correlation is specifically very high (r = 0.61; p= 0.003) when the FE near the start codon is the strongest (Figure 4E). The intermediate sequences are expected to have a stronger effect on initiation when this process is less efficient (i.e. when it is more rate limiting). Thus, according to our model we expect to see stronger correlation between protein levels and the number of intermediate sequences when the mRNA folding in the region surrounding the START codon is strong (Figure 4F.2).

When calculating the partial Spearman correlation between the number of sub-sequences that interact in a *weak* manner with the rRNA and the PA of the GFP variants, the correlation is negative and significant (r=-0.32; p= 8.5·10^−5^). This further validates our conjecture that translation efficiency in this case is indeed related to interactions that are neither very strong, nor very weak or absent. It also suggests that this effect on translation efficiency is related to the pre-initiation step and not the elongation step, otherwise we would expect positive correlation with weak interaction (Supplementary section S.6 and Figure S5).

We also analyzed *E. coli* genes by their mRNA half-life(33) to assess how selection for intermediate interactions varies among them. We found that genes with shorter half-life tend to have more intermediate interactions (Supplementary section S.7). It is possible that these genes undergo stronger selection to include intermediate interactions since their corresponding mRNAs ‘have less time’ to initiate translation. Thus, the reported results discussed here suggest that the diffusion of the small ribosomal sub-unit is probably relatively fast.

It is known that mRNAs tend to localize in certain regions in the cell(34), meaning that if we can keep the ribosome close to a certain mRNA we also keep it close to other mRNA’s. If a certain mRNA ‘captures’ a ribosome, then undergoes degradation this ribosome will likely remain close to other nearby mRNAs. It is also possible that due to compartmentalization and aggregation of many mRNA molecules the interaction with the small sub-unit of one mRNA can be ‘helpful’ for a nearby mRNA.

Finally, we created a computational biophysical model that describes the movement of the small ribosomal sub-unit along the transcript. In this model the movement is influenced by the intermediate interactions (Figures 4G and 4H). The model indicates that adding intermediate interaction along the transcript improves the initiation rate and termination rate even if the intermediate sequence is near the 3’ end of the gene. It also demonstrates the advantage of intermediate interactions over weak or strong ones in most of the transcript as intermediate interactions in the transcript optimize the translation rate. We conclude that intermediate rRNA-mRNA interactions along the transcript enhance small ribosomal sub-unit diffusion to the start codon with resultant improvements in the translation rate (Supplementary S.8).

## Discussion

This study revolutionizes current understandings of how mRNA-rRNA interactions affect translation efficacy and efficiency throughout all stages of translation in many prokaryotes. We provide multiple streams of evidence that in many bacterial phyla the 16S rRNA plays a role in regulating all stages of translation via its interaction with the mRNA (Supplementary S.9 and figures S7 and S8). Such interactions are usually directly correlated with organism growth rates (Supplementary S.10 and figures S9 and S10). Evidence for the effect of these interactions on translation does not appear in organisms without an aSD sequence in the rRNA (Supplementary section S.11 and figures S11-S16).

Previous work has identified pieces of this puzzle, such as the importance of the aSD-SD interactions for translation initiation(1, 2), and some initial evidences that these interactions may be deleterious in the coding regions(19, 35). This study is novel in expanding our understanding of the effect of such interactions *throughout* the translation process - from pre-initiation small sub-unit diffusion, through initiation, elongation, and termination. It presents new evidence of selection for intermediate interactions in the transcript, and for strong aSD-SD interactions upstream the stop codons and downstream the start codon with much details about the distributions of the reported patterns in transcripts. This study also provides novel insights into how evolutionary forces reflect trade-offs and non-linear relationships between the effect and fitness derived from different interaction strengths in different parts of the mRNA transcript, and how these vary across levels of gene expression and phyla. In addition, we show how the reported patterns correlated and/or affected by other features of the transcripts such as the strength of the mRNA folding near the start codon, the mRNA stability, the aSD patterns in the rRNA, etc. The results are supported by various scientific approaches including selection detection, molecular biology experiments, and biophysical modeling.

Our findings also shed light on the biophysics of translation in bacteria and the conformation of the rRNA of the small ribosomal subunit and its interaction with mRNA molecules during the various steps of the translation process. The interactions described herein can also be implemented in engineered transcripts for efficient expression in various bacterial species. Rigorous molecular evolution models should take into account our findings that even within coding regions, the selection of nt and codons that comprise mRNAs are shaped by rRNA-mRNA interactions and the way in which these affect translation efficiency.

Our results demonstrate various complex trade-offs and non-monotonous relations between the optimal rRNA-mRNA interaction strength across transcripts. For example, *increasing* the rRNA-Mrna interaction strength inside coding regions, both *decreases* elongation efficiency and *increases* the initiation efficiency. Thus, evolution in these regions tends to shape coding regions such that they include intermediate (not too strong, not too weak) levels of rRNA-mRNA interactions. At the 3’ end of the coding region, strong rRNA-mRNA interactions tend to *improve* termination fidelity, although they *decrease* translation rates and may increase ribosomal traffic jams. Thus, evolution at the 3’ end tends to shape the end of coding regions such that they include more rRNA-mRNA interactions when they do not decrease translation efficiency (i.e. in genes that are less translationally efficient). Further research should examine these non-monotonous relations in high resolution both from the molecular evolution, engineering, and biophysical points of view.

## Materials and Methods

### The analyzed organisms

We analyzed 551 bacteria genome from the following phyla or classes: Alphaproteobacteria, Betaproteobacteria, Cyanobacteria, Deltaproteobacteria, Gammaproteobacteria, Gram positive bacteria, Purple bacteria, Spirochaetes bacteria. In addition, we analyzed 76 bacteria genome across the tree of life that do not have a canonical aSD sequence in their 16S rRNA. Finally, we analyzed 196 bacteria genome with known growth rates. The full lists of analyzed organisms can be found in Table S1, Table S2, Table S3. All of the bacterial genomes were downloaded from the NCBI database (https://www.ncbi.nlm.nih.gov/) on October 2017. In addition to the coding regions, for each gene, we also analyzed the 50nt upstream of the start codon and the 50nt downstream of the stop codon (approximating the end of the 5’UTR, and the beginning of the 3’UTR respectively).

### Protein levels

*E.coli*endogenous protein abundance data was downloaded from PaxDB (http://pax-db.org/download), we used “*E. coli* – whole organism, EmPAI” published in 2012.

### The rRNA-mRNA strength prediction

The prediction of rRNA-mRNA interaction strength is based on the hybridization free energy between two sub-sequences: The first sequence is a sequence from the mRNA and the second sequence is the aSD from the rRNA. This energy was computed based on the Vienna package RNAcoFold(36), which computes a common secondary structure of two RNA molecules. Lower, more negative free energy is related to stronger hybridization.

We assumed that the interacting sub-sequence at the 16S rRNA 3’ end is *UCCUCC* (3’ to 5’). However, when we remove this assumption (change the aSD sequence) and infer it in an unsupervised manner, the results remain similar.

### The rRNA-mRNA interaction strength profiles and selection strength

The rRNA-mRNA interaction strength profiles include the predicted rRNA-mRNA hybridization strength for each position in each transcript (UTRs and coding regions), and in each bacterium. We analyzed the average profile of each bacterium.

We calculated the interaction strength between all 6 nucleotide sequences along each transcript (UTR’s and coding sequences) with the 16S rRNA aSD. For each possible genomic position along the transcripts we performed a statistical test (empirical P-value) to decide if the potential rRNA-mRNA interaction in this position is significantly strong, intermediate, or weak. In order to decide if a position (across the entire transcriptome) tends to include sub-sequences with certain rRNA-mRNA interaction strength (strong, intermediate or weak) we compared it to the properties of sub-sequences observed in a null model in the same position (see further details regarding the null model below). We also created Z-score maps of the strength of interactions, see Supplementary section S.12 and Figure S17.

### The null model

For each bacterial genome we designed 100 null model mRNA randomizations. UTR regions were generated with nucleotide permutation, preserving the nucleotide distribution, including the GC content of the original mRNA. Coding regions were generated by permuting synonymous codons while preserving codon frequencies, amino acid order and content; and GC content of the original mRNA.

Similar rRNA-mRNA interaction strength profiles as the ones described above were computed for the randomized versions of the transcripts, to compute p-values related to possible selection for strong/intermediate/weak rRNA-mRNA interactions.

We computed an empirical p-value for every position in the trancriptome of a certain organism. To this end, the average rRNA-mRNA interaction strength in the position was compared to the average obtained in all of the randomized genomes. The p-value was computed based on the number of times the real genome average was higher or lower (depend on the hypothesis we checked) than the null model average. A significant position is a position with a p-value smaller than 0.05.

For more details on the validity of the null models see Supplementary sections S.13 and S.14 and figures S18 and S19.

### The intermediate rRNA-mRNA interaction definition

In order to define intermediate interaction strength, we devised an unsupervised adaptive optimization model that infers the thresholds (upper and lower) of intermediate interaction strength. We assumed that coding regions tend to include intermediate interactions and chose the thresholds that maximize their number in comparison to the null model. Specifically, we implemented an algorithm that in each iteration change the selected upper and lower thresholds mentioned above such that the number of significant positions in terms of the number of intermediate interactions in the real genome compared to the null model will be maximal.

The first iteration thresholds were selected as follows; we created a distribution histogram of interaction strength in the region with the strong canonical SD interaction in the 5’UTR of each bacterium (positions -8 through -17, Figure 1B). We calculated the area under the strong interaction distribution. We initially chose the ‘high’ (strongest interaction strength -- more negative free energy) and ‘low’ (weakest interaction strength -- less negative free energy) thresholds to be the interaction strength such that the area up to the chosen threshold interaction value was 5% of the total distribution area from each side of the curve.

To study the properties of the selected thresholds, we created the interaction strength histograms for two regions in the 5’UTR (Figure 4A): 1) The distribution of strong interaction strength as mentioned above. 2) The distribution of interaction strength in positions -40 to -50 at the 5’UTR upstream of the START codon (where we do not expect to see strong rRNA-mRNA interaction, as this region doesn’t have a known role in translation initiation).

Next, we looked at the positions of the two inferred thresholds in comparison to these two histograms; as can be seen in Figure 4A, they tend to appear in the region between the two historgams supporting the hypothesis that these are indeed intermediate interaction strength.

To further quantitatively validate the inferred thresholds, we calculated the area under the two histograms mentioned above induced by the two inferred thresholds. The ratio between these two areas (the first one divided by the second one) was computed: A ratio larger than one suggests that it is more probable that the inferred thresholds are related to (intermediate) interactions between the rRNA and mRNA than to lack of interactions; indeed, in most bacteria (503/551) the ratio was larger than one (Figure 4D).

### Relation between the number of intermediate rRNA-mRNA interactions in the coding regions and heterologous protein levels

We aimed at showing that intermediate sequences in the coding region of a gene directly improve its translation initiation efficiency, and thus its protein levels. Hence, we calculated the partial Spearman correlations between the number of intermediate interaction sequences in the GFP variant and the heterologous protein levels (PA), based on 146 synonymous GFP variants that were expressed from the same promoter and the same UTR(28).

The control variables were the CAI (Codon Adaptation Index – a measure of codon bias) and folding energy (FE) near the start codon. We defined an area of intermediate interactions according to the thresholds received by our model in *E. coli* and we expanded it by 20% to allow maximum intermediate interactions in this synthetic system (which is expected to differ from endogenous genes). The correlation was indeed positive and significant (r=0.35; P=2·10^−5^), suggesting that variants with more sub-sequences in the coding region that bind to the rRNA with an intermediate interaction strength tend to have higher PA.

### Ribosome Profiling

*E. coli* ribosome footprint reads were obtained from(26) (SRR2340141,3-4). *E. coli* transcript sequences were obtained from NCBI (NC_000913.3). Sequenced reads were mapped as described in (37) with the following minor modifications. We trimmed 3’ adaptors from the reads using Cutadapt (38) (version 1.17), and utilized Bowtie (39) (version 1.2.1) to map them to the *E. coli* transcriptome. In the first phase, we discarded reads that mapped to rRNA and tRNA sequences with Bowtie parameters ‘–n 2 –seedlen 21 –k 1 --norc’. In the second phase, we mapped the remaining reads to the transcriptome with Bowtie parameters ‘–v 2 –a --strata --best --norc –m 200’. We filtered out reads longer than 30nt and shorter than 23nt. Unique alignments were first assigned to the ribosome occupancy profiles. For multiple alignments, the best alignments in terms of number of mismatches were kept. Then, multiple aligned reads were distributed between locations according to the distribution of unique ribosomal reads in the respective surrounding regions. To this end, a 100nt window was used to compute the read count density (total read counts in the window divided by length, based on unique reads) in the vicinity of the *M* multiple aligned positions in the transcriptome, and the fraction of a read assigned to each position was 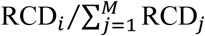. The location of the A-site was set for each read length by the peak of read distribution upstream of the stop codon for that length.

After creating the ribosome profiling distributions, for each gene, we calculated the number of positions with strong rRNA-mRNA interactions in the last 20 nucleotides of the coding region (the location of the reported signal, Figure 3A). We ranked the genes according to their ‘number of strong positions’, and defined the 10% highest/lowest ranking genes. For the highest and lowest ranking genes, we calculated the average Ribo-seq read count in the first 20 nucleotides of the 3’ UTR (the closest region to the stop codon), Figure 3E.

### Z-score calculation in highly and lowly expressed genes

To validate the reported signals, we performed all of our analyses on highly and lowly expressed genes of *E. coli*. We chose the highly and lowly expressed genes according to their PA (20% highest and lowest PA values), and computed Z-scores as explained in the next sub-sections.

#### Highly vs. lowly: Selection for Strong rRNA-mRNA interactions at the 5’UTR end and at the beginning of the coding region

We calculated the Z score based on the rRNA-mRNA interaction strength in all possible positions in the 5’UTR and coding region in the highly and lowly expressed genes.

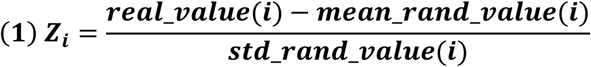

- *Z*_*i*_ - Z-score in position *i*.
- *real_value(i)* – rRNA-mRNA interaction strength in position *i*.
- *mean_rand_value(i)* – Average rRNA-mRNA interaction strength in position i in all of the randomizations.
- *std_rand_value(i* – Standard deviation of rRNA-mRNA interaction strength in position *j* in all of the randomizations.

From a statistical point of view, we defined each gene by two values according to the reported signal: 1) Minimum Z-score value in position -8 through -17 in the 5’UTR. 2) Minimum Z-score value in position 1 through 5 at the beginning of the coding region. The regions were selected according to the reported signal in Figure 1B.

We performed two Wilcoxon rank-sum tests to estimate the p-values for the two reported signals in highly vs. lowly expressed genes.

#### Highly vs. lowly: Selection against strong rRNA-mRNA interactions at the beginning of the coding sequence

We calculated the Z-score (as described above) based on the rRNA-mRNA interaction strength of each position in the first 400nt of the coding region in the highly and lowly expressed genes.

We performed Wilcoxon rank-sum tests to estimate the p-values of the reported signals.

#### Highly vs. lowly: Z-score calculation of selection for strong mRNA-rRNA interactions at the end of the coding sequence

In this case, we calculated the Z-score (as described above) based on the rRNA-mRNA interaction strength of each position in the last 20nt of the coding region in each bacterium.

For each bacterium, we found the position with a minimum Z-score value (strongest interaction compared to the null model). We created a histogram of the positions of the strongest Z-scores in the last 20nt of the coding region (Figure 3C), and a histogram based on gene expression levels (Figure 3D).

### Selection against strong interaction in the coding region in positions that are not upstream to a close AUG codon

To detect signal of selection for/against strong interaction in the coding region after excluding positions that are upstream to a close start codon we preformed the following analysis. We considered the *E*. coli genomes (both real and randomized versions) and in each gene we “marked”, position that are up to 14 positions upstream of an AUG (in all frames). We then computed p-value related to selection for strong rRNA-mRNA interactions (as mentioned before) but when we consider only the non-marked positions (both in the real and the randomized genomes). The result can be seen in Figure S2.

### Selection against strong interaction at the end of the coding region – read-through experiment

#### Plasmids construction

We used plasmid pRX80 and modified it by deleting the lac I repressor gene and the CAT selectable marker. The resulting plasmid contained the RFP and GFP genes in tandem, both are expressed from a promoter with two consecutive lac operator domains. The plasmid contains also the pBR322 origin of replication and the Kanamycin resistance gene as a selectable marker. Because the 2 operator sequences caused instability at the promoter region, we replaced the promoter region with a lacUV promoter with only one operator sequence. The resulting plasmid, pRCK28 was now used for the generation of variants which differ in the 40 last nucleotides of the RFP ORF. The variants include synonymous changes composed of both ribosome binding site at 3 energy ranges and which also alter the local folding energy (LFE) of the 40 last nucleotides of the RFP ORF end. The variable sequences where synthesized as G-blocks and Gibson assembly was used to replace the relevant region of the pRCK28 plasmid, generating 9 variants as described in Figure S4 (A, C). The resulting variable plasmids were transformed into competent *E. coli* DH5 cells. Colonies were selected on LB Kanamycin plates. A few candidates were PCRed and sequenced to verify the synonymous changes in each variant.

#### Fluorescent Tests

Single colonies of each variant as well as of the original pRCK28 clone and of a negative control (an *E. coli* clone harboring a Kanamycin resistant plasmid at the same size of pRC28 but without any fluorescent gene) were grown overnight in LB-Kanamycin. Cells were then diluted and 10,000 cells were inoculated into 110ul defined medium (1X M9 salts, 1mM thiamine hydrochloride, 2% glucose, 0.2% casamino acids, 2mM MgSO_4,_ 0.1mM CaCl_2_) in 96 well plates. For each variant 2 biological repeats and 4 technical repeats of each were used. A fluorimeter (Spark-Tecan) was used to run growth and fluorescence kinetics. For growth, OD at 600 nm data were collected. For red fluorescence, excitation at 555nm and emission at 584nm were used. For green fluorescence, excitation at 485nm and emission at 535nm were used. Data was analyzed and normalized by subtracting the auto fluorescence values of the negative control, and by calculating the fluorescence to growth intensity ratios.

#### Western blot analyses

Cells were grown overnight, 1 ml cultures were concentrated by centrifugation and lysed using the BioGold lysis buffer supplemented with lysozyme. Total protein lysates were resolved on Tris glycin 4-15% acrylamide mini protein TGX stain free gels (BioRad). Proteins were transferred to nitrocellulose membranes using the trans-blot Turbo apparatus and transfer pack. Membranes were incubated in blocking buffer (TBS+1% casein) for 1 hr at room temprature. Anti GFP and/or anti RFP antibodies (Biolegend) were used at 1:5K, for 1 hr in blocking buffer, at room temperature to probe the GFP and RFP expression. Goat anti-mouse 2^nd^ antibody was then applied at 1:10K dilution. ECL was used to generate a binding signal.

### Unified biophysical translation model of the reported signals

We developed a computational simulative model of translation that includes the pre-initiation, initiation and elongation phases. Our model is based on a mean field approximation of the TASEP model(40). All of the model parameters are based on rRNA-mRNA interaction strength.

The model consists of two types of ‘particles’: 1. Small sub-units of the ribosome (pre-initiation): in this case, detachment/attachment and bi-direction movement of the particles is possible along the entire transcript. 2. Ribosome (elongation): the movement is unidirectional (from the 5’ to the 3’ of the mRNA) and possible only in the coding region; the initiation rate is affected by the density of the small sub-units of the ribosome at the ribosomal binding site (RBS) (Supplementary section S.8 and Figure S6).

## Acknowledgment

This study was supported in part by a fellowship from the Edmond J. Safra Center for Bioinformatics at Tel-Aviv University, by the Ofakim research fellowship by Miri and Efraim and The Koret-UC Berkeley-Tel Aviv University Initiative in Computational Biology and Bioinformatics. We thank Hadas Zur for helpful comments

## REFERENCES

1. Shine J, Dalgarno L. 1975. Determinant of cistron specificity in bacterial ribosomes. Nature 254:34–38.

2. Kozak M. 1999. Initiation of translation in prokaryotes and eukaryotes. Gene 234:187–208.

3. Nakagawa S, Niimura Y, Miura K -i., Gojobori T. 2010. Dynamic evolution of translation initiation mechanisms in prokaryotes. Proc Natl Acad Sci 107:6382–6387.

4. Shine J, Dalgarno L. 1974. The 3’-Terminal Sequence of Escherichia coli 16S Ribosomal RNA: Complementarity to Nonsense Triplets and Ribosome Binding Sites. Proc Natl Acad Sci 71:1342–1346.

5. Dontsova O, Kopylov A, Brimacombe R. 1991. The location of mRNA in the ribosomal 30S initiation complex; site-directed cross-linking of mRNA analogues carrying several photo-reactive labels simultaneously on either side of the AUG start codon. EMBO J 10:2613–20.

6. Dahlberg AE, Brimacombe R, Atmadja J, Stiege W, Schuler D, Crick FHC, Cundliffe E, Denman R, Negre D, Cunningham PR, Nurse K, Colgan J, Weitzmann C, Ofengand J, Hall CC, Johnson D, Cooperman BS, Hausner T-P, Atmadja J, Nierhaus KH, Herr W, Chapman NM, Noller HF, Hui AS, DeBoer HA, Hui AS, Eaton DH, Boer HA de, Jacob WF, Santer M, Dahlberg AE, Meier N, Goringer HU, Kleuvers B, Scheibe U, Eberle J, Szymkowiak C, Zacharias M, Wagner R, Melancon P, Lemieux C, Brakier-Gingras L, Moazed D, Noller HF, Moazed D, Noller HF, Moazed D, Noller HF, Moazed D, Stolk BJ Van, Douthwaite S, Noller HF, Moazed D, Robertson JM, Noller HF, Murgola EJ, Hijazi KA, Goringer HU, Dahlberg AE, Noller HF, Asire M, Barta A, Douthwaite S, Goldstein T, Gutell R, Moazed D, Normanly J, Prince JB, Stern S, Triman K, Turner S, Stolk B Van, Wheaton V, Weiser B, Woese CR, Noller HF, Stern S, Moazed D, Powers T, Svensson P, Changchien L-M, Nomura M, Held WA, Poldermans B, Bakker H, Knippenberg PH Van, Prince JB, Taylor BH, Thurlow DL, Ofengand J, Zimmermann RA, Santer M, Sigmund C, Ettayebi E, Morgan EA, Spirin AS, Steiner G, Kuechler E, Barta A, Tapprich WE, Hill WE, Tapprich WE, Goss DJ, Dahlberg AE, Thomas CL, Gregory RJ, Winslow G, Muto A, Zimmermann RA, Thompson JF, Hearst JE, Thompson J, Cundliffe E, Dahlberg AE, Trifonov EN, Vester B, Garrett RA, Weiss RB, Dunn DM, Atkins JF, Gesteland RF, Weiss RB, Dunn DM, Dahlberg AE, Atkins JF, Gesteland RF, Woese CR, Wool I, Zwieb C, Jemiolo DK, Jacob WF, Wagner R, Dahlberg AE. 1989. The functional role of ribosomal RNA in protein synthesis. Cell 57:525–529.

7. Steitz J a, Jakes K. 1975. How ribosomes select initiator regions in mRNA: base pair formation between the 3’ terminus of 16S rRNA and the mRNA during initiation of protein synthesis in Escherichia coli. Proc Natl Acad Sci U S A 72:4734–4738.

8. Shaham G, Tuller T. 2018. Genome scale analysis of Escherichia coli with a comprehensive prokaryotic sequence-based biophysical model of translation initiation and elongation. DNA Res 25:195–205.

9. Li GW, Oh E, Weissman JS. 2012. The anti-Shine-Dalgarno sequence drives translational pausing and codon choice in bacteria. Nature 484:538–541.

10. Liu X, Jiang H, Gu Z, Roberts JW. 2013. High-resolution view of bacteriophage lambda gene expression by ribosome profiling. Proc Natl Acad Sci 110:11928–11933.

11. Subramaniam AR, DeLoughery A, Bradshaw N, Chen Y, O’Shea E, Losick R, Chai Y. 2013. A serine sensor for multicellularity in a bacterium. Elife 2:e01501.

12. Schrader JM, Zhou B, Li G-W, Lasker K, Childers WS, Williams B, Long T, Crosson S, McAdams HH, Weissman JS, Shapiro L. 2014. The Coding and Noncoding Architecture of the Caulobacter crescentus Genome. PLoS Genet 10:e1004463.

13. Diwan GD, Agashe D. 2016. The frequency of internal shine-dalgarno-like motifs in prokaryotes. Genome Biol Evol 8:1722–1733.

14. Vasquez KA, Hatridge TA, Curtis NC, Contreras LM. 2016. Slowing Translation between Protein Domains by Increasing Affinity between mRNAs and the Ribosomal Anti-Shine-Dalgarno Sequence Improves Solubility. ACS Synth Biol 5:133–145.

15. Hockenberry AJ, Jewett MC, Amaral LAN, Wilke CO. 2018. Within-gene shine-dalgarno sequences are not selected for function. Mol Biol Evol 35:2487–2498.

16. Tuller T, Carmi A, Vestsigian K, Navon S, Dorfan Y, Zaborske J, Pan T, Dahan O, Furman I, Pilpel Y. 2010. An evolutionarily conserved mechanism for controlling the efficiency of protein translation. Cell 141:344–354.

17. Tuller T, Zur H. 2015. Multiple roles of the coding sequence 5’ end in gene expression regulation. Nucleic Acids Res 43:13–28.

18. Fredrick K, Ibba M. 2010. How the sequence of a gene can tune its translation. Cell 141:227–229.

19. Yang C, Hockenberry AJ, Jewett MC. 2016. Depletion of Shine-Dalgarno Sequences Within Bacterial Coding Regions Is Expression Dependent 6:3467–3474.

20. Weiner I, Shahar N, Marco P, Yacoby I, Tuller T. 2019. Solving the Riddle of the Evolution of Shine-Dalgarno Based Translation in Chloroplasts. Mol Biol Evol 36:2854–2860.

21. Scharff LB, Childs L, Walther D, Bock R. 2011. Local absence of secondary structure permits translation of mrnas that lack ribosome-binding sites. PLoS Genet 7.

22. Zur H, Tuller T. 2013. New Universal Rules of Eukaryotic Translation Initiation Fidelity. PLoS Comput Biol 9.

23. Bonetti B, Fu L, Moon J, Bedwell DM. 1995. The Efficiency of Translation Termination is Determined by a Synergistic Interplay Between Upstream and Downstream Sequences inSaccharomyces cerevisiae. J Mol Biol 251:334–345.

24. Namy O, Rousset J-P, Napthine S, Brierley I. 2004. Reprogrammed Genetic Decoding in Cellular Gene Expression. Mol Cell 13:157–168.

25. Dos Reis M, Wernisch L. 2009. Estimating translational selection in eukaryotic genomes. Mol Biol Evol 26:451–461.

26. Mohammad F, Woolstenhulme CJ, Green R, Buskirk AR. 2016. Clarifying the Translational Pausing Landscape in Bacteria by Ribosome Profiling. Cell Rep 14:686–694.

27. Peeri M, Tuller T. 2020. High resolution modeling of the selection on local mRNA folding strength in coding sequences across the tree of life. Genome Biol.

28. Kudla G, Murray AW, Tollervey D, Plotkin JB. 2009. Coding-sequence determinants of gene expression in Escherichia coli. Science (80-) 324:255.

29. Sharpl PM, Li W. 1987. The codon adaptation index - a measure of directional synonymous codon usage bias, and its potential applications. Nucleic Acids Res 15:1281–1295.

30. Hall MN, Gabay J, Débarbouillé M, Schwartz M. 1982. A role for mRNA secondary structure in the control of translation initiation. Nature 295:616–618.

31. Tuller T, Waldman YY, Kupiec M, Ruppin E. 2010. Translation efficiency is determined by both codon bias and folding energy. Proc Natl Acad Sci 107:3645–3650.

32. Nackley AG, Shabalina SA, Tchivileva IE, Satterfield K, Korchynskyi O, Makarov SS, Maixner W, Diatchenko L. 2006. Human catechol-O-methyltransferase haplotypes modulate protein expression by altering mRNA secondary structure. Science 314:1930–3.

33. Bernstein JA, Khodursky AB, Lin P-H, Lin-Chao S, Cohen SN. 2002. Global analysis of mRNA decay and abundance in Escherichia coli at single-gene resolution using two-color fluorescent DNA microarrays. Proc Natl Acad Sci 99:9697–9702.

34. Nevo-Dinur K, Nussbaum-Shochat A, Ben-Yehuda S, Amster-Choder O. 2011. Translation-independent localization of mRNA in E. coli. Science (80-) 331:1081–1084.

35. Hockenberry AJ, Amaral AN, Jewett MC, Wilke CO. 2018. Selection removes Shine-Dalgarno-like sequences from within protein coding genes Assessing the conservation status of Shine-Dalgarno-like sequence.

36. Hofacker IL. Vienna RNA secondary structure server.

37. Diament A, Tuller T. 2016. Estimation of ribosome profiling performance and reproducibility at various levels of resolution. Biol Direct 11:24.

38. Martin M. 2011. Cutadapt removes adapter sequences from high-throughput sequencing reads. EMBnet.journal 17:10–12.

39. Langmead B, Trapnell C, Pop M, Salzberg SL. 2009. Ultrafast and memory-efficient alignment of short DNA sequences to the human genome. Genome Biol 10:R25.

40. Reuveni S, Meilijson I, Kupiec M, Ruppin E, Tuller T. 2011. Genome-Scale Analysis of Translation Elongation with a Ribosome Flow Model. PLoS Comput Biol 7:e1002127.

